# Molecular Dynamics Reveals the Effects of Temperature on Critical SARS-CoV-2 Proteins

**DOI:** 10.1101/2021.01.24.427990

**Authors:** Paul Morgan, Chih-Wen Shu

## Abstract

Severe Acute Respiratory Syndrome Coronavirus 2 (SARS-CoV-2) is a newly identified RNA virus that causes the serious infection Coronavirus Disease 2019 (COVID-19). The incidence of COVID-19 is still increasing worldwide despite the summer heat and cool winter. However, little is known about seasonal stability of SARS-CoV-2. Herein, we employ Molecular Dynamics (MD) simulations to explore the effect of temperature on four critical SARS-CoV-2 proteins. Our work demonstrates that the spike Receptor Binding Domain (RBD), Main protease (Mpro), and nonstructural protein 3 (macro X) possesses extreme thermos-stability when subjected to temperature variations rendering them attractive drug targets. Furthermore, our findings suggest that these four proteins are well adapted to habitable temperatures on earth and are largely insensitive to cold and warm climates. Furthermore, we report that the critical residues in SARS-CoV-2 RBD were less responsive to temperature variations as compared to the critical residues in SARS-CoV. As such, extreme summer and winter climates, and the transition between the two seasons, are expected to have a negligible effect on the stability of SARS-CoV-2 which will marginally suppress transmission rates until effective therapeutics are available world-wide.

## BACKGROUND

Severe Acute Respiratory Syndrome Coronavirus 2 (SARS-CoV-2) is the virus responsible for the global spread of the respiratory illness Coronavirus Disease 2019 (COVID-19).^1^ On March 11, 2020, World Health Organization (WHO) publicly declared the SARS-CoV-2 outbreak a pandemic.^2^ As of January 24, 2021, over 100 million cases have been confirmed worldwide, resulting in over two million deaths. This global public health crisis began during the peak of winter in the norther hemisphere where the average temperature during the winter months from December to March, averages at 273K (3°C). Several vaccine candidates have now been approved and are currently being distributed world-wide.^3^ At present, there are very few reports that investigate the stability of SARS-CoV-2 in response to changes in temperature.

The SARS-CoV-2 genome is comprised of nearly 30,000 nucleotides. These nucleotides encodes four structural proteins including the spike (S) protein, membrane (M) protein, envelope (E) protein, and nucleocapsid (N) protein.^4,5^ The envelope glycoprotein spike (S) attaches to host cells during infection through binding with Angiotensin-converting enzyme 2 (ACE-2).^6^ The spike glycoprotein is synthesized as a precursor and cleaved into two subunits, including S1 and S2. The S1 contains the RBD, which has critical amino acids required for interaction with cellular ACE-2.^6^

Of the four proteins, Mpro is one of the most attractive viral targets for antiviral drug discovery against SARS-CoV-2 receiving major attention during the first SARS-CoV outbreak in 2004.^5^ Viral proteases have proven to be well-validated drug targets against chronic infections caused by the hepatitis C virus (HCV) and human immunodeficiency virus (HIV).^5^ Mpro plays a crucial role during viral replication by cleaving overlapping polyproteins to mature functional proteins.^5^ Consequently, inhibiting Mpro can stall the production of viral particles and therefore suppress the symptoms and severity of the COVID-19 infection.

The macro X domain is one of the most novel proteins of SARS-CoV-2 encoded by nonstructural protein 3 (nsp3).^7^ Nsp3 is a large multidomain membrane-bound protein, and its clearest role in viral replication is cleaving the rep polyprotein.^7^ Although the biological role of ADP-ribose binding is still poorly understood, macro X domain is also a promising drug target.^7^

Of all the structural proteins, the N protein is a highly immunogenic and abundantly expressed during infection.^7^ Although the SARS–CoV-2 S protein is currently being used as a leading target antigen in vaccine development the complex mechanisms of viral entry lends itself to complications in vaccine response.^7^ In contrast to the SARS–CoV-2 S protein, the N gene is more conserved and stable, with 90% amino acid homology and fewer mutations over time, also making it an ideal drug target.^2,7^

To date, the rate of infection worldwide has unarguably demonstrated that SARS–CoV-2 is quite stable when subjected to cold and warm temperatures.^8^ However, the effects of temperature on the RBD, Mpro, macro X, and the nucleocapsid protein remains unclear and requires urgent clarity with respect to their potential as stable drug targets.

## METHODS

SARS-CoV-2 RBD (PDB: 6VW1), Mpro (PDB: 6M03), macro X (PDB: 6YWL), nucleocapsid (PDB: 6M3M) and SARS-CoV RDB (PDB: 2AJF) were cleaned, side chains were fixed, and the structures were minimized. CMIP titration was performed and then the system was solvated in transferable intermolecular potential with 3 points (TIP3P) water molecules and ions were added to equalize the total system charge. The steepest decent algorithm was used for initial energy minimization until the system converged at Fmax 500 kJ/ (mol · nm). Water and ions were allowed to equilibrate around the protein in two phases. The first phase of equilibration was at a constant number of particles, volume, and temperature (NVT) while the second phase was at a constant number of particles, pressure, and temperature (NPT). The system was allowed to equilibrate at a reference temperature of 300 K, or reference pressure of 1 bar for 2.5 ps at a time step of 2 fs. The production simulations were performed for 50 ns with 2 fs time intervals at 255K(−18°C), 261K(−12°C), 266K(−7°C), 272K(−1°C), 277K(3.85°C°C), 283K(10°C), 288K(15°C), 294K(15°C), 300K(27°C), 305K((32°C), 310K((37°C), 322K(49°C), 333K(60°C), 344K(71°C), 355K(82°C), 366K(93°C), 377k(104°C). RMSD were calculated for the RBD, Mpro, macro X, and nucleocapsid. While RMSF was extracted for the subRBD which is comprised of the critical residues responsible for the initial interaction with ACE-2. RMSF was also determined for randomly selected residues throughout the RBD from the N-terminus to the C-terminus for standardization. Average structures at each temperature were generated as Protein Data Bank (PDB) files and Audio Video Interleave (AVI) files for simulation playback. Data analysis and correlation plots were generated using SPSS, Prism 5 and Python.

## RESULTS AND DISCUSSION

### Relating RMSD and protein stability

Herein, we employ Molecular Dynamics (MD) simulations to assess the effect of temperature on the SARS-CoV-2 RBD, Mpro, macro X and the nucleocapsid. We observed that subjecting the proteins to changes in temperature, from 255K(−18°C) to 322K(49°C), which corresponds to the transition from the peak of winter to the peak of summer. The videos for MD simulation of these four proteins at temperature 255K and 322K were provided as supporting information. We found that these four proteins had predominantly negligible effects as shown on the Root Mean Square Deviation (RMSD)(Fig. S1-S4). RMSDs of Cα-atoms of a protein as a function of simulation time give an estimate of the stability of protein structure. Macro X was demonstrated to be the most thermo-stable protein with very negligible changes in average RMSD across the temperature series (Fig. 1). The nucleocapsid on the other hand, was the least thermos-stable protein with an average RMSD of 2 Å across the temperature series. The RBD and Mpro had comparable stability averaging 0.5 Å in RMSD (Fig. 1).

**Fig. 1.**
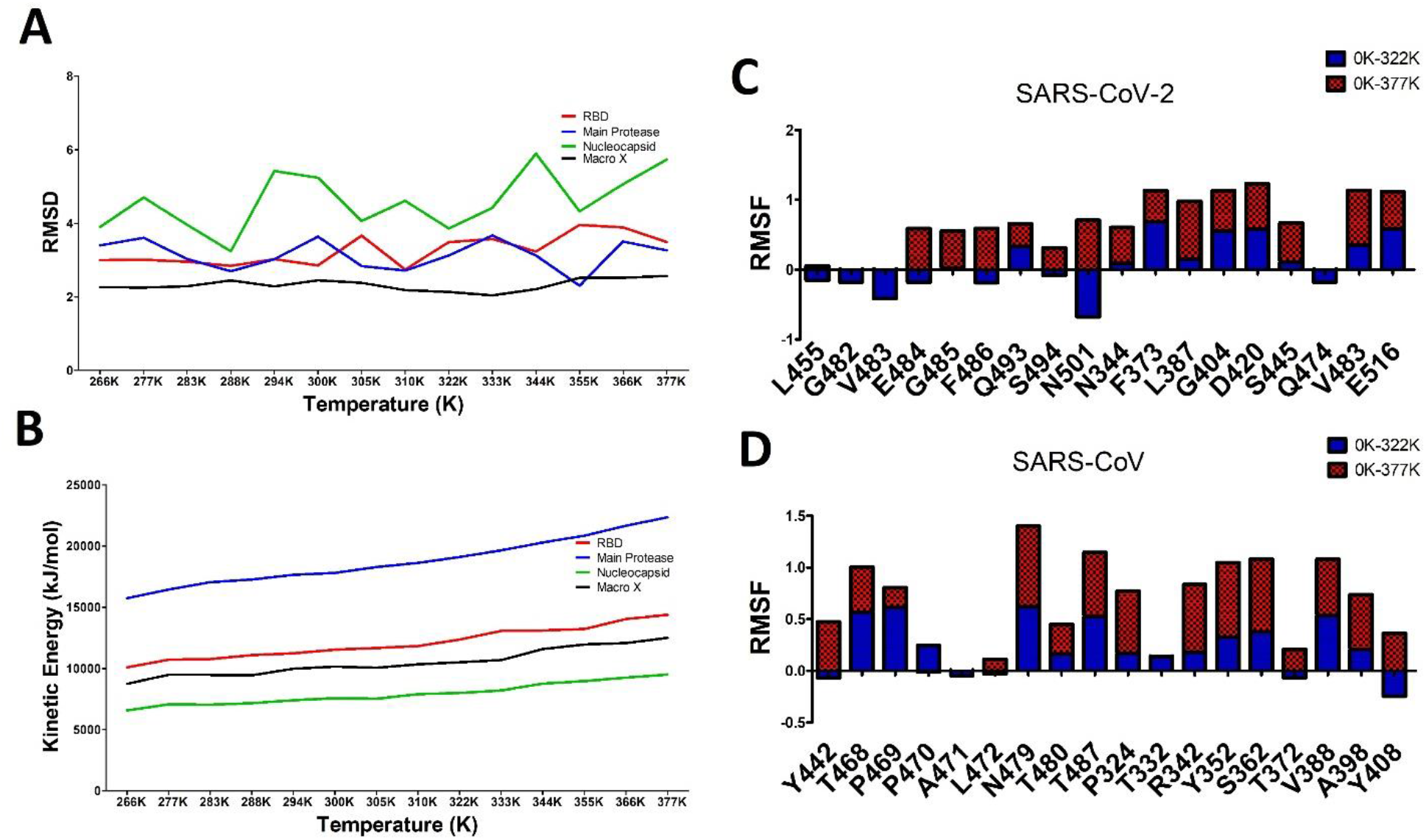
(A) Shows the average RMSD for the RBD, Mpro, Macro X, and the nucleocapsid for a total simulation time of 50 ns across a temperature series from 255K to 377K. (B) Depicts the corresponding average kinetic energy of the globular proteins during the simulation. The RBD, Mpro, and Macro X demonstrated the greatest thermo-stability across the temperature series. Shows the RMSF over 50 ns for several critical residues in the (C) RBD of SARS-CoV-2 (D) RBD of SARS-CoV across a temperature gradient from 0K to 377K. The critical residues in the SARS-CoV-2 sub-RBD appear to be more resistant to changes in temperature as compared to SARS-CoV. L455, G482, V483, S494 and Q474 of SARS-CoV-2 are highly thermo-stable and may play a critical role in efficient host recognition and infection.

It is also worthwhile noting that the nucleocapsid demonstrated the lowest average kinetic energy (KE) across the temperature series while Mpro had the highest average KE. Macro X and the RBD had comparable KE and were the least responsive when subjected to the temperature series. Our findings suggest that the RBD, Mpro, and Maco X, are inherently thermos-stable, rendering them ideal drug targets with potentially desirable drug binding kinetics. This is because secondary structural changes are often a consequence of inhibitor binding. As such, monitoring variations in secondary structures as a function of simulation time is instrumental in determining the formation of a stable protein-inhibitor complex.

### RMSF of the critical residues in the RBD

Next, we investigated the effect of temperature on the Root Mean Square Fluctuation (RMSF) of the critical residues in the RBD. These residues are ultimately responsible for initializing the interaction with ACE-2, the entry point for the infection of pneumocytes in the lungs. We report that the critical residues in SARS-CoV-2 RBD were less responsive to temperature variations as compared to the critical residues in SARS-CoV (Fig. 2). It is worthwhile noting that the critical residues are predominantly located on the subRBD of both strains. This is consistent with the understanding that sequence regions involved in binding tend to show significant interspecific, temperature-related differences in flexibility.^9^

**Fig. 2.**
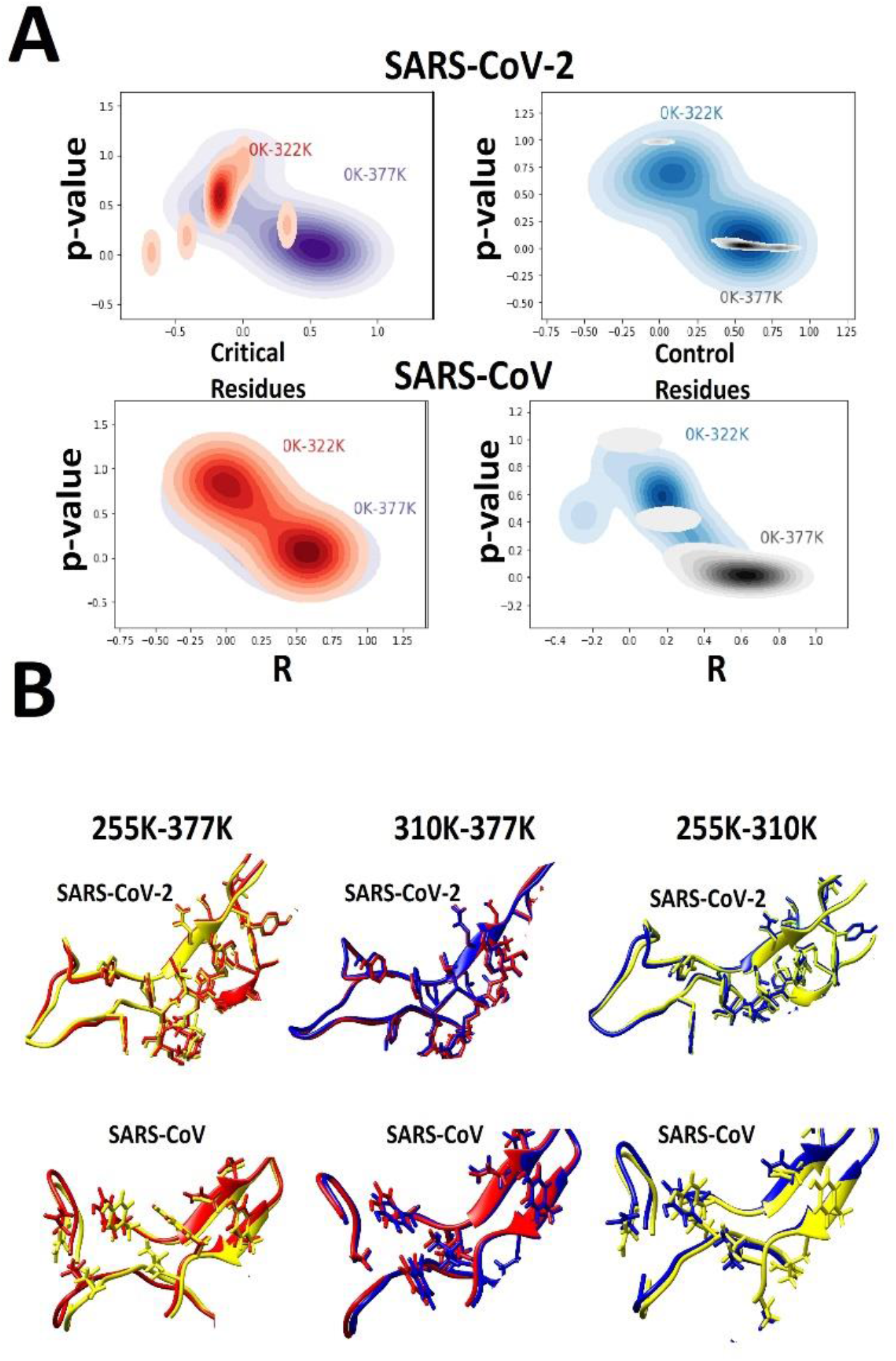
(A) Pearson’s Correlation Coefficient (R) to RMSD for both SARS-CoV-2 and SARS-CoV RBD from 255K to 322K and 255K to 377K. 2D Kernel Density Estimates (KDE) of R and p-value of RMSD for SARS-CoV-2 and SARS-CoV critical and control residues. Depicts the sensitivity of SARS-CoV-2 and SARS-CoV critical and control residues from 255K to 377K. (B) Superimposition of the average structure of the subRBD at 255K (yellow), 310K (blue) and 377K (red) for SARS-CoV-2 and SARS-CoV.

We further discriminated amongst the critical residues and inferred that L455, G482, and V483 had negligible responses to the change in temperature from 255K to 377K.^2^ This provides further insight into the individual roles of the of critical residues with respect to the enhanced rigidity, and thermal stability of the subRBD.^2^

This increased degree of rigidity in the SARS-CoV-2 subRBD, and, still yet, enigmatically enhanced flexibility of the global RBD is perhaps a strategic evolutionary mutation that can at minimum, partly explain the increased affinity and likely increased specificity for ACE-2. Conventionally, it is understood that flexibility is positively correlated with specificity.^9–12^ Whereas, increased rigidity is correlated with increased affinity.^12^ Flexibility and affinity are generally regarded as inverse correlates.^9,11,12^ Thus, the elevated rigidity and reduced flexibility in the SARS-CoV-2 subRBD might be the driving force behind the significantly enhanced increase in affinity for ACE-2 (Fig. 3). Likewise, the overall increase in globular flexibility of SARS-CoV-2 RBD may have far-reaching implications with regards to its specificity for ACE-2 (Fig. 3).

**Fig. 3.**
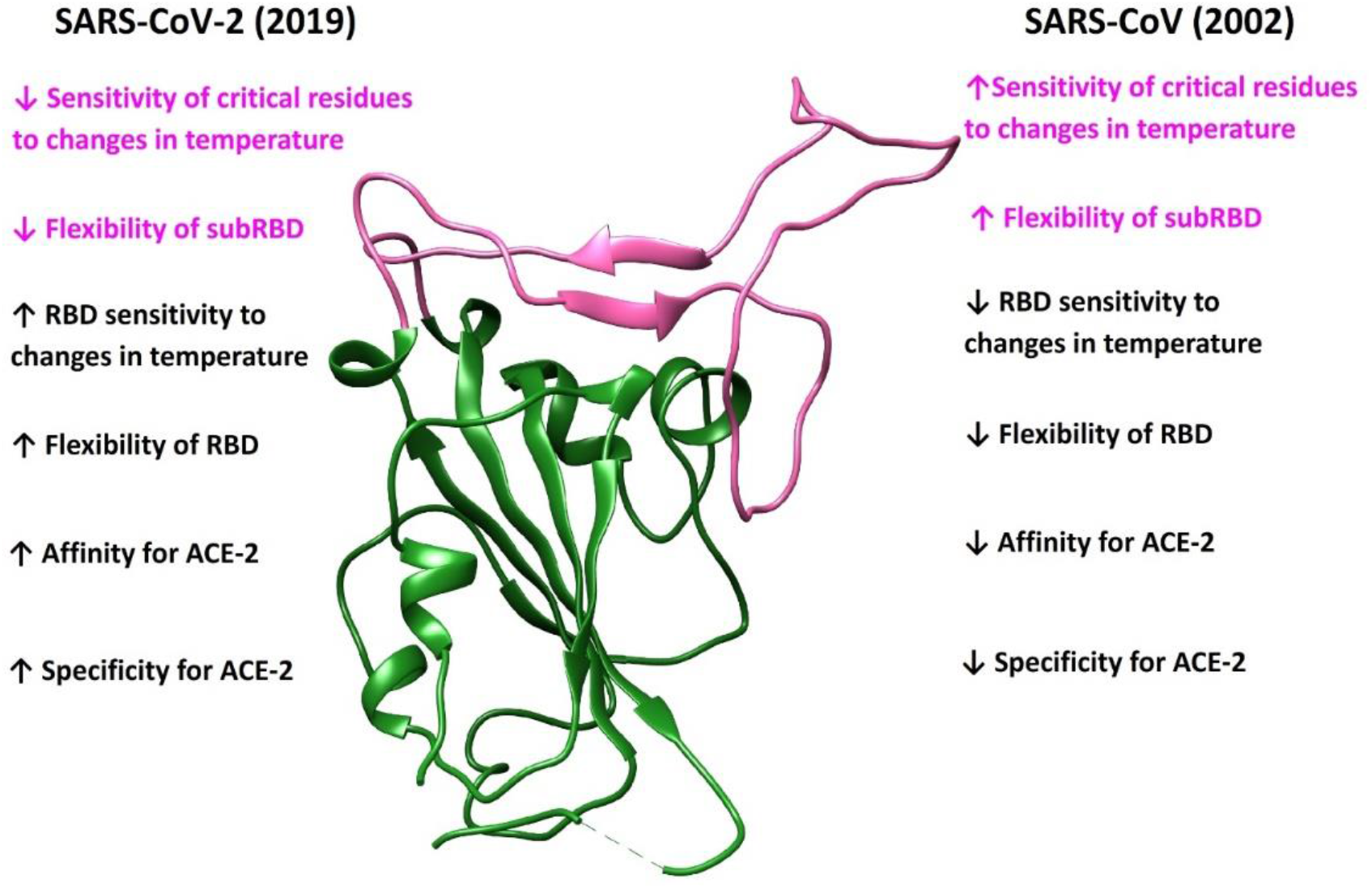
Proposed evolution of SARS-CoV (2002) to SARS-CoV-2 (2019). These strategic residue mutations have resulted in a RBD that more versatile, more efficient and more resistant to temperature variations. This implication of this evolution is evidenced by the high transmitability of SARS-CoV-2 as compared to its predecessor SARS-CoV.

Nevertheless, there are practical limitations that must be considered as well. We have only computationally evaluated the thermo-stability of four critical SARS-CoV-2 proteins in response to temperature changes within an aqueous environment. However, this is not a comprehensive assessment of the plentiful proteins and conditions that exist as SARS-CoV-2 lingers within our communities. For instance, how is the stability of the RBD affected by surfaces such as cloth, wood and metal?^13^ Furthermore, it is not a trivial task to attempt simulating changes in humidity using MD simulations, despite its intimate relationship with temperature. Also, our MD simulations did not take into account the complex N-linked glycan at reside N-343 found on the core RBD.^14^ The shielding of receptor binding sites by glycans is a common feature of viral glycoproteins, which is consistent with that observed in SARS-CoV.^14^

Last, we have only considered the RBD, Mpro, macro X, and the nucleocapsid, which is but a small fraction of the entire pathogen. MD simulations are also currently incapable of simulating the homeostatic responses of the internal viral proteins and their complex interactions when subjected to extreme environmental parameters like changes in pH, pressure, and humidity along with temperature variations. Finally, the coronavirus machinery is intriguingly complex and may have other stabilizing mechanisms and protein-protein interactions to cope with environmental stresses that is brought forth during the transition between summer and fall, fall and winter, winter and spring, and spring and summer.

## CONCLUSION

In conclusion, the evolution of SARS-CoV to SARS-CoV-2 suggests a complex evolutionary, advantageous interplay that has resulted in this profound global pandemic (Fig 3.). Furthermore, the implications of our study suggests that four essential SARS-CoV-2 proteins are well adapted to habitable temperatures on earth and exhibit extreme thermo-stability.^15^ As a consequence, we have thus far observed a negligible effect on the transmitability of SARS-CoV-2 with respect to changes in temperature between the winter months and summer months. More importantly, our study establishes a framework for elucidating stable promising drug targets for SARS-CoV-2.

## ETHICS STATEMENT

We have confirmed this is an original work and has not been published elsewhere. The authorship is limited to those who have made a significant contribution to the study as follows: PM and CWS investigated, interpreted the data, and wrote the manuscript. PM prepared the figures, performed the statistical analyses in the manuscript.

## ACKNOWLEDGMENT

The work was supported by the Ministry of Science and Technology MOST (108-2320-B-110 - 008 −MY3) for corresponding author Chih-Wen Shu. This version of the manuscript is in the preprint.

