## Supplemental Figures1-4 for "Molecular Dynamics Reveals the Effects of Temperature on Critical SARS-CoV-2 Proteins"

**SUPPLEMENTARY**

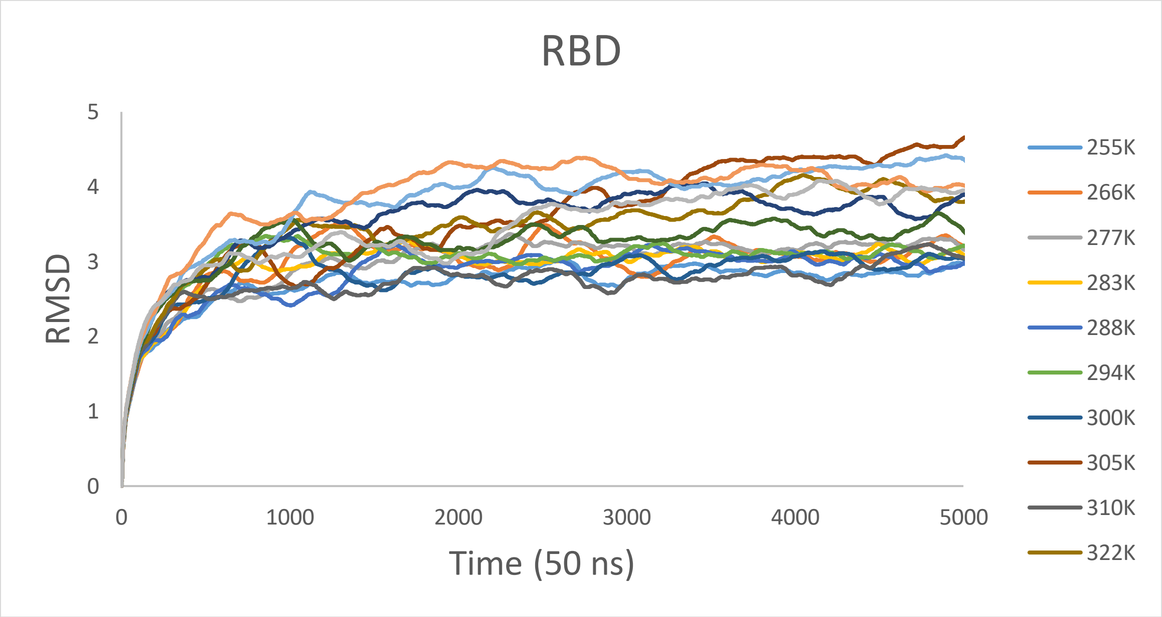


**Fig S1.** Shows the RMSD for the RBD of SARS-CoV-2 for 50 ns across a temperature series from 255K to 377K.

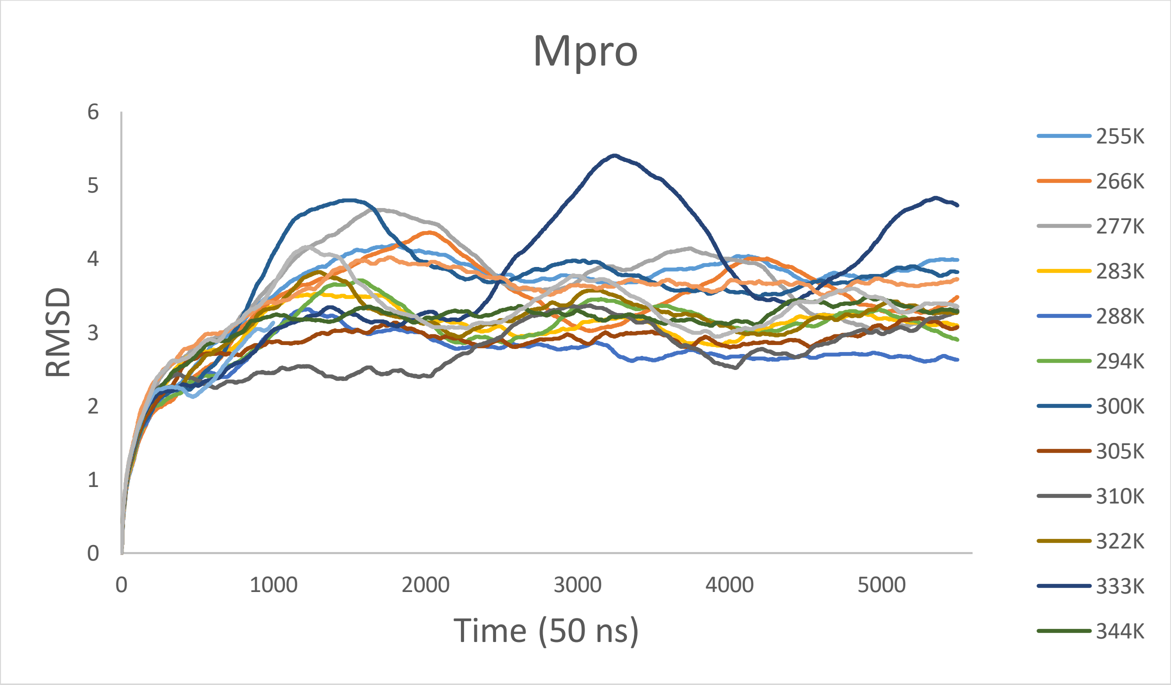


**Fig S2.** Shows the RMSD for the Mpro of SARS-CoV-2 for 50 ns across a temperature series from 255K to 377K.

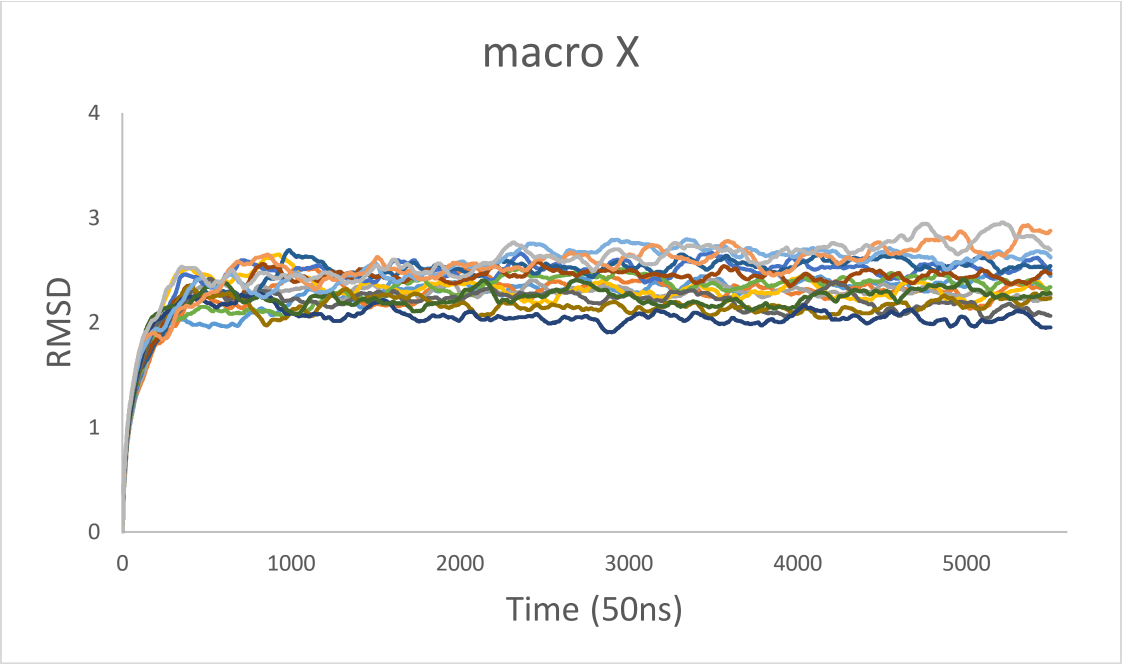


**Fig S3.** Shows the RMSD for macro X of SARS-CoV-2 for 50 ns across a temperature series from 255K to 377K.

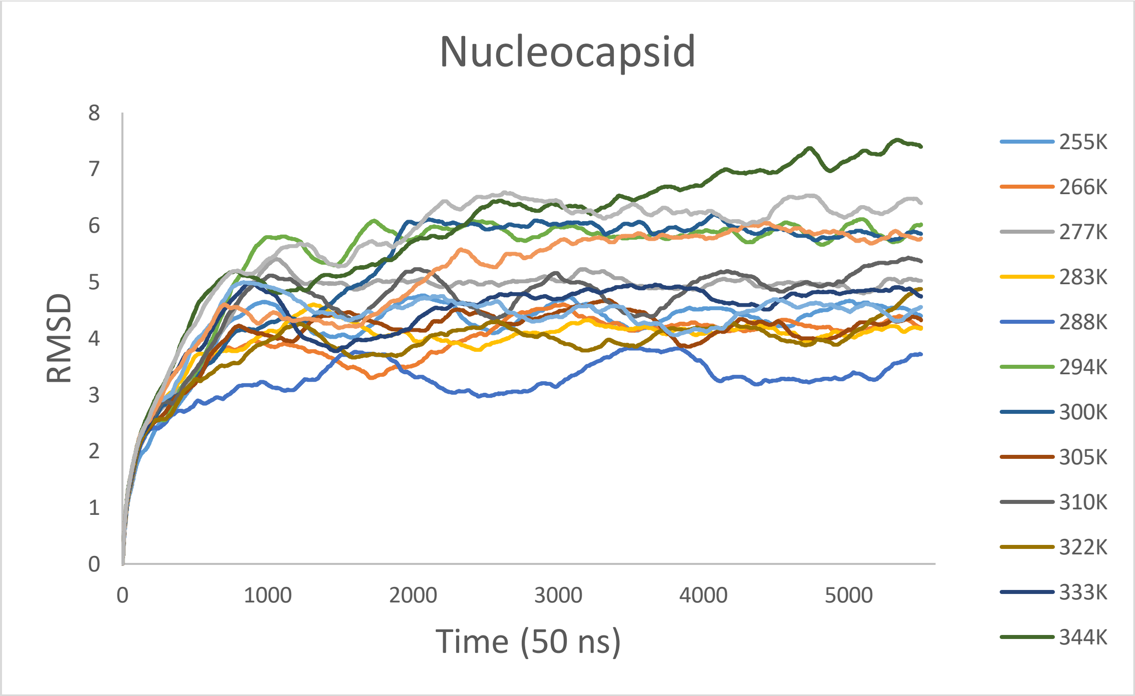


**Fig S4.** Shows the RMSD for the nucleocapsid of SARS-CoV-2 for 50 ns across a temperature series from 255K to 377K.
